# Two distinct neuronal populations in the rat parafascicular nucleus oppositely encode the engagement in stimulus-driven reward-seeking

**DOI:** 10.1101/2021.02.14.431166

**Authors:** Mehdi Sicre, Julie Meffre, Frederic Ambroggi

## Abstract

The thalamus is a phylogenetically well-preserved structure. Known to densely contact cortical regions, its role in the transmission of sensory information to the striatal complex has been widely reconsidered in recent years. The parafascicular of the thalamus (Pf) has been implicated the orientation of attention towards salient sensory stimuli. In a stimulus-driven reward seeking task, we sought to characterize the electrophysiological activity of Pf neurons in rats. We observed a predominance of excitatory responses over inhibitory responses for all events of the task. Neurons responded more strongly to the stimulus compared to lever-pressing and collecting reward, confirming the strong involvement of the Pf in sensory information processing. The use of long sessions allowed us to compare neuronal responses to stimuli when the animal engaged in action or when it did not. We distinguished two populations of neurons responding in an opposite way: MOTIV+ neurons responded more intensively to stimuli followed by a behavioral response than those that did not. Conversely, MOTIV-neurons responded more strongly when the stimulus was ignored by the animal. In addition, MOTIV-neurons excitations appeared at a shorter latency after the stimulus than MOTIV+ neurons. Through this encoding, Pf could perform an early selection of environmental stimuli transmitted to the striatum according to motivational level.

**HIGHLIGHTS:**

- Pf neurons respond to reward-predicting stimuli and reward-related actions
- MOTIV+ Pf neurons were more active to stimuli evoking reward-seeking
- MOTIV- Pf neurons were more active to stimuli ignored by the animal
- Stimuli-evoked excitations latencies were shorter in MOTIV- than MOTIV+ neurons

## INTRODUCTION

The historic view of the thalamus as an ensemble of nuclei relaying sensory information to the cortex (Ariens-Kappers et al., 1936) has largely been reconsidered. The thalamus is a phylogenetically ancient structure that evolved before the expansion of the neocortex (Butler, 1994). Thus, if thalamic nuclei unequivocally provides essential information to the cerebral cortex through theirs projections, it is not surprising that they also intensively connects subcortical regions (James et al., 2011; Rikhye et al., 2018). In particular, the associative thalamic nuclei from the midline and intralaminar group send strong projections to the striatal complex (Van der Werf et al., 2002). This structure is considered the entry station of the basal ganglia, a subcortical network participating in the selection and the control of voluntary actions (Redgrave et al., 1999; Hikosaka et al., 2014; Sirigu and Duhamel, 2016). This pattern of connectivity has challenged a key element of the classical model of the basal ganglia stating that the input signal to the striatum originates from the cerebral cortex (Alexander, 1986). Some authors even radically proposed that the thalamus, rather than the cortex, may be the dominant source of sensory information to the basal ganglia (McHaffie et al., 2005; Redgrave and Gurney, 2006; Redgrave et al., 2011). This appreciation of thalamic afferences to the striatum has largely incentivized their investigation at the cellular and network level (Ding et al., 2010; English et al., 2012; Doig et al., 2014; Mandelbaum et al., 2019) and several studies have now begun to address their functional implications at the behavioral level (Matsumoto et al., 2001; Hamlin et al., 2009; Bradfield et al., 2013; Do-Monte et al., 2017; Díaz-Hernández et al., 2018; Meffre et al., 2019).

In particular, the primate center médian (CM)/parafascicular nucleus (Pf) complex from the caudal intralaminar nuclei has attracted a lot of attention. The CM is not present in the rodent brain but it has been largely documented that the primate CM/Pf complex is equivalent to the rodent Pf (Jones and Leavitt, 1974; Smith et al., 2004, 2011). An extensive topographic projection follows a medio-lateral axis in the Pf to a ventromedial-dorsolateral axis in the striatal complex in both rats (Berendse and Groenewegen, 1990, 1991; Van der Werf et al., 2002) and mice (Mandelbaum et al., 2019). The medial Pf projects to the ventral region of the striatum (i.e. nucleus accumbens, NAc) while the lateral Pf projects to the dorsolateral striatum. A similar topography has also been described in primates (Jones and Leavitt, 1974; Groenewegen and Berendse, 1994; Smith et al., 2004).

Electrophysiological recordings have shown that primate CM/Pf neurons are rapidly excited by stimuli from a variety of sensory modalities (auditory, visual and somatosensory). These activations depend on the temporal unpredictability of stimuli and their association with rewards (Matsumoto et al., 2001), suggesting that the CM/Pf is involved in attentional orientating toward salient sensory stimuli (Minamimoto and Kimura, 2002; Kimura et al., 2004). Furthermore, it has been reported that CM/Pf neurons are more excited by stimuli predicting a small than a large reward (Minamimoto et al., 2005) and that the magnitude of these excitations inversely correlate with reaction times (Minamimoto et al., 2014).

Thus, the CM/Pf could provide important information to the striatum to adequately control the engagement in action in response to temporally unexpected stimuli predictive of rewards. To our knowledge, the response of Pf neurons to reward-predictive stimuli has not been investigated in rodents. The aim of this study was to fill this gap by recording single neuron activity in the Pf in a task where rats had the opportunity to engage in reward-seeking by performing an instrumental action in response to a predictive stimulus. We used long sessions to compare Pf neuronal responses to stimuli when rats decided to engage in reward-seeking and when they did not. We found a population of neurons that responded more intensively to reward-predicting stimuli the animals responded to compared to those they ignored. A second population, excited to stimuli at shorter latency, had an opposite pattern of activity. We propose that these two populations may respectively promote and refrain the engagement in action through their connections to the NAc.

## EXPERIMENTAL PROCEDURES

### Subjects

Experiments were conducted on male Long-Evans rats (Charles Rivers, France) weighing ~300g on arrival. Rats were immediately housed individually on a 12 h light/dark cycle. Experiments were conducted during the light phase. After one week of habituation, rats were placed under food restriction; food rations were adjusted daily to maintain the body weight at ~90% of their free-feeding body weight. All experiments were performed in accordance with the guidelines on animal care and use of the European guidelines (European Community Council Directive, 2010/63/UE) and National guidelines.

### Training

All experiments were conducted in operant chambers containing two house lights, a tone speaker, a retractable lever and a reward receptacle located on one wall of the chamber (Med Associates, Vermont, USA). Liquid sucrose (10%) was delivered as a reward in the receptacle by a syringe pump. During the first 2 days, rats were trained to obtain 50μl of sucrose by spontaneously entering into the reward receptacle. When 300 rewards per hour were obtained, rats were run on a fixed-ratio 1 (FR1) schedule with a 10 s-time-out: the lever was constantly extended in the chamber and a lever-press triggered the delivery of 50μl of sucrose into the receptacle. When rats reached the criterion >100 lever-presses per hour, they were advanced to a stimulus-driven reward-seeking FR1 task.

### Stimulus-driven reward-seeking task

Rats were run daily on the task for 3 hours. The stimulus was composed of the extension of the lever associated with a 300 ms white noise (85 dB). When the stimulus was presented, rats had 10 s to press the lever to obtain the reward. Each trial was followed by a variable interval schedule averaging 45 s (from 30 to 60 s). If the rats did not respond on the lever within 10 s, the lever retracted and the intertrial interval was re-initialized. Surgery was performed when rats reached the criterion of >80% responses during the first hour of the session.

### Surgery

Rats were anesthetized with isoflurane (5%, Tem Sega, France) and placed in a stereotaxic apparatus (Kopf Instruments, California, USA). Anesthesia level was then adjusted with 0.5-2 % isoflurane during the maintenance phase. Before skin incision, a subcutaneous injection of lidocaine (1mg/ml, Lurocaine MedVet,) was performed. Bundles of 8 electrodes were attached to custom-made-microdrive devices (du Hoffmann et al., 2011) that allowed to lower the electrode bundles by 80 μm increments. Electrodes were implanted in the Pf bilaterally for 5 animals and unilaterally in 2 animals (one in the left and the other in the right hemisphere) at the following stereotaxic coordinates: AP: −4.1, ML: +/-1, DV: −5.4 mm relative to the Bregma. The microdrive and the connector were secured to the skull with bone screws, adhesive cement (C&B Metabond, Phymep, France) and dental acrylic (Phymep, France). After surgery, a prophylactic analgesic treatment (Buprenorphine, 0.05 mg/kg, Vetergesic, France) was administered, and rats were given at least 7 days of recovery with *ad libitum* access to food. After recovery, rats were placed in food restriction and re-trained until reaching previous criteria.

### Electrophysiology

#### Recording procedure

Electrophysiological recordings were conducted as described previously (Sicre et al., 2020) during the entire 3 hour-long sessions. Animals were connected to the electrophysiological acquisition system (SpikeGadget LLC, California, USA). The 32-channel headstage streamed data at 30kHz per channel and was connected to a low-torque HDMI commutator that allowed the animals to be free of their movement in the chamber. Between sessions, electrode bundles were lowered by 80 or 160 μm to record news set of neurons. Unfiltered data were transferred from the data acquisition main control unit to a data acquisition computer where it was visualized and saved. Digitally-filtered data (0.2-6kHz) was used for spike sorting.

#### Spike sorting

Recorded data were analyzed offline with OfflineSorter (Plexon Inc, Texas, USA) to isolate spikes from single neurons with principal component analysis. Inter-spike interval distributions, cross-correlograms and auto-correlograms were used to ensure that the activity of single neurons was isolated. Only well isolated waveforms with characteristics that were constant over the entire recording session were included in this study. Sorted units were exported to NeuroExplorer 4.135 (Nex Technologies, Colorado, USA) and Matlab R2018b (MatWorks Inc, Massachusetts, USA) for further analysis.

#### Electrophysiological analyses

##### Waveform Analysis

In our data set, 290 neurons showed waveforms with a negative followed by a positive deflection. The remaining 95 neurons displayed the opposite pattern. The spike width was assessed by the time elapsed between the first and second extremum independently from the sign of the first and second deflections. The spike width of 14 neurons could not be determined due to technical issues and we then excluded from the analyses involving this parameter.

##### Response detection

Peri-stimulus time histograms (PSTHs) were constructed with smoothed (lowess method, span=4) 20ms- and 2ms-time-bins. PSTHs constructed around the behavioral events (stimulus presentation, lever-press and reward collection) were used to detect excitations and inhibitions and the time at which they occurred. The 10 s period before the presentation of the stimulus was used as a baseline period. Excitation and inhibition to each event was determined by the presence of at least 3 consecutive bins above the 99 % (for excitations) or below the 1 % (for inhibitions) confidence interval of the baseline during the analysis windows (0 to 250ms after the stimulus, −1000 to 500ms around lever-presses and reward delivery). Onset was determined by the time of the first of 5 consecutive bins falling outside the confidence interval. The offset was determined in analogy, by searching the first of 10 consecutive bins within the confidence interval.

##### Deconvolution

To isolate the activity of temporally close events, we used a deconvolution method as described previously (Ghazizadeh et al., 2010; Ambroggi et al., 2011). Briefly, the model assumes that the total firing rate of a neuron in each trial is equal to the linear sum of the contributions of each event-related firing, delayed by the event latencies in that trial. Here, we deconvolved single event responses for each neuron using the optimal number of iterations that had a cross validation error lower than the PSTHs.

##### Data normalization and plotting

Color-coded maps and average PSTHs across neurons were constructed with 20ms- and 2ms-bins. Prior to averaging, the firing rate of each neuron during each bin was z-score-transformed: (F_i_ - F_mean_)/F_sd_ where F_i_ is the firing rate of the i^th^ bin of the PSTH, and F_mean_ and F_sd_ are, respectively, the mean and the SD of the firing rate during the 10 s preceding stimulus onset.

### Statistical analyses

We sought to compared the trials to which the animal engaged in reward-seeking with those it did not. Thus, we selected the sessions containing at least 20 trials of each type to conduct reliable analysis of the neuronal data. For behavioral analyses, the primary dependent variables were the percentage of responding to the stimulus and the latency to lever-press after the presentation of the stimulus.

For electrophysiological data, the primary dependent variables were the baseline firing rate, the maximal frequency reached (measured by averaging the top 5% of instantaneous frequencies during the baseline period), the coefficient of variation (measured during the baseline period), the spike width, the proportion of responsive neurons, the mean z-score normalized firing (0-100 ms post-stimulus and −500 to 1000 ms around lever-presses and reward collection), the onset latency and the durations of the responses. These variables were analyzed with paired, unpaired t-tests or ANOVAs. When appropriate, a Bonferroni test was used to conduct post-hoc comparisons. Proportions were analyzed with χ^2^ tests and distributions with Kolmogorov-Smirnov tests. All results were considered significant at p < 0.05.

### Histology

Animals were deeply anesthetized with a solution of pentobarbital (Euthasol, 140 mg/kg). A local anesthesia was performed on the thorax in the incision site (Lurocaine 10 mg/kg). Each electrodes site was labeled by passing a DC current of 20 μA for 7 s. Rats were perfused intracardially with phosphate buffered saline following by 10 % formalin solution. Brains where removed, post-fixed in 10 % formalin and placed in 30 % sucrose for 3 days. Brains were sectioned at 40 μm on a cryostat and slices were stained with cresyl violet. Reconstruction of the recording sites was made based on the final location of the electrodes.

## RESULTS

### Behavioral analysis

Rats were run in a stimulus-driven FR1 task where a 300ms-auditory stimulus was presented unexpectedly (in average every 45 s) in conjunction with the extension of a lever lasting up to 10 s. A lever-press triggered the immediate delivery of a 10 % sucrose reward in an adjacent receptacle and the retraction of the lever (Fig. 1A, B). We sought to analyze neuronal activity when the rats attended to the stimulus and when they did not. In order to obtain a sufficient number of trials of each type, we ran 3-hour long sessions. For all rats, the percentage of lever-pressing in response to the reward-predictive stimulus and the latency were variable across sessions (5 to 9 sessions per rat for 7 rats, total of 46 sessions Fig. 1C). Overall, the mean percentage of responding was 60.9 ± 2.8 % with an average stimulus-lever-press latency of 2.64 ± 0.14 s.

**Fig.1.**
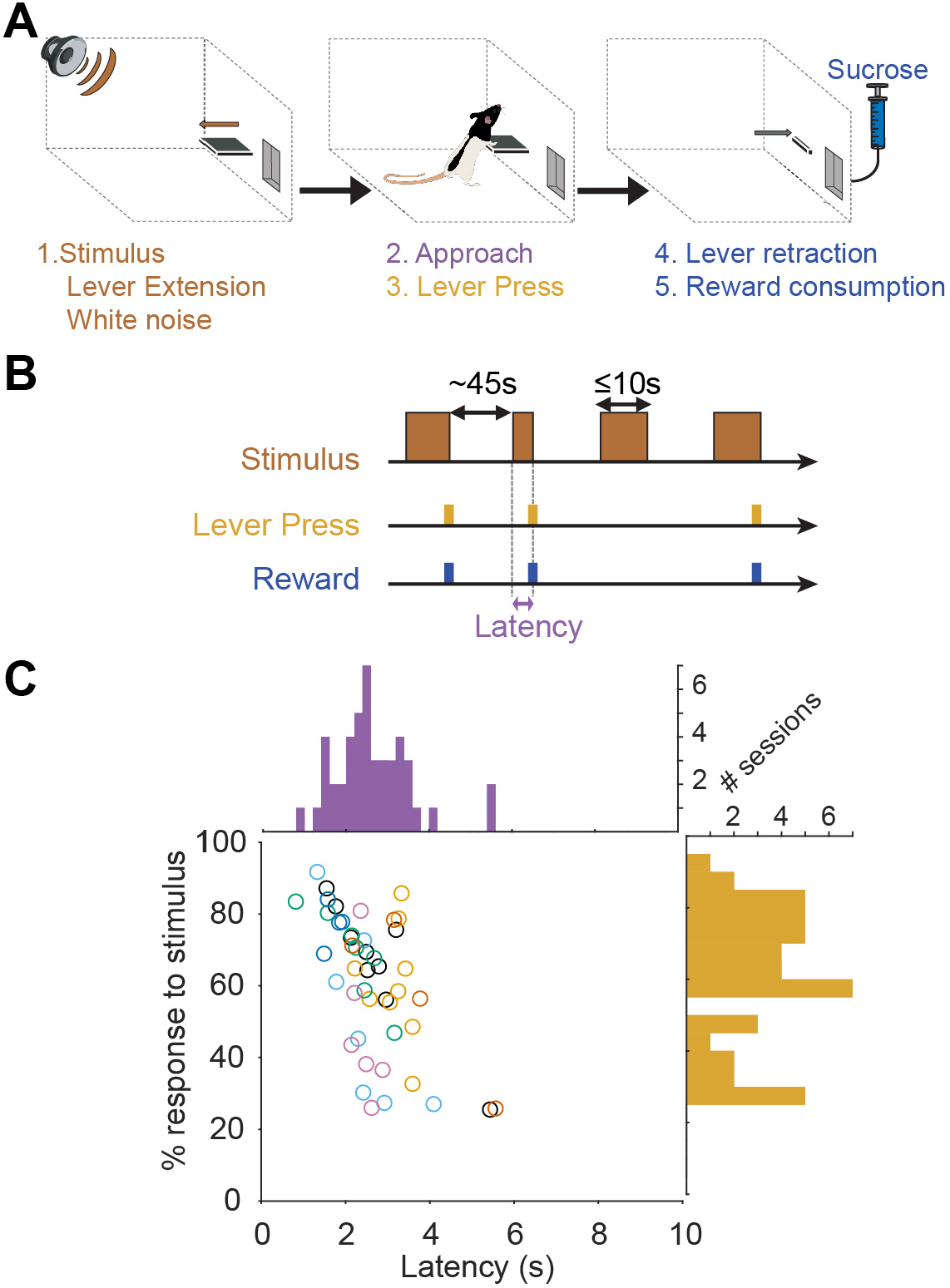
Behavioral performance in the stimulus-driven reward seeking task. **A.** Task diagram showing the sequence of events in one trial. **B.** Temporal structure of the task. **C.** Individual percentage of responses and latencies for all sessions (n=46). Animals (n=7) are color-coded. Histograms represent the distributions of the percentage of responses (yellow) and latencies (purple).

### Basic electrophysiological properties of Pf neurons

We recorded single unit activity of 399 Pf neurons (see Fig. 2 for histological reconstruction of the electrode sites). We first conducted analyses of neuronal responses to the different events on the trials the animals responded to the stimuli by lever-pressing and collecting the rewards (Fig. 3).

**Fig.2.**
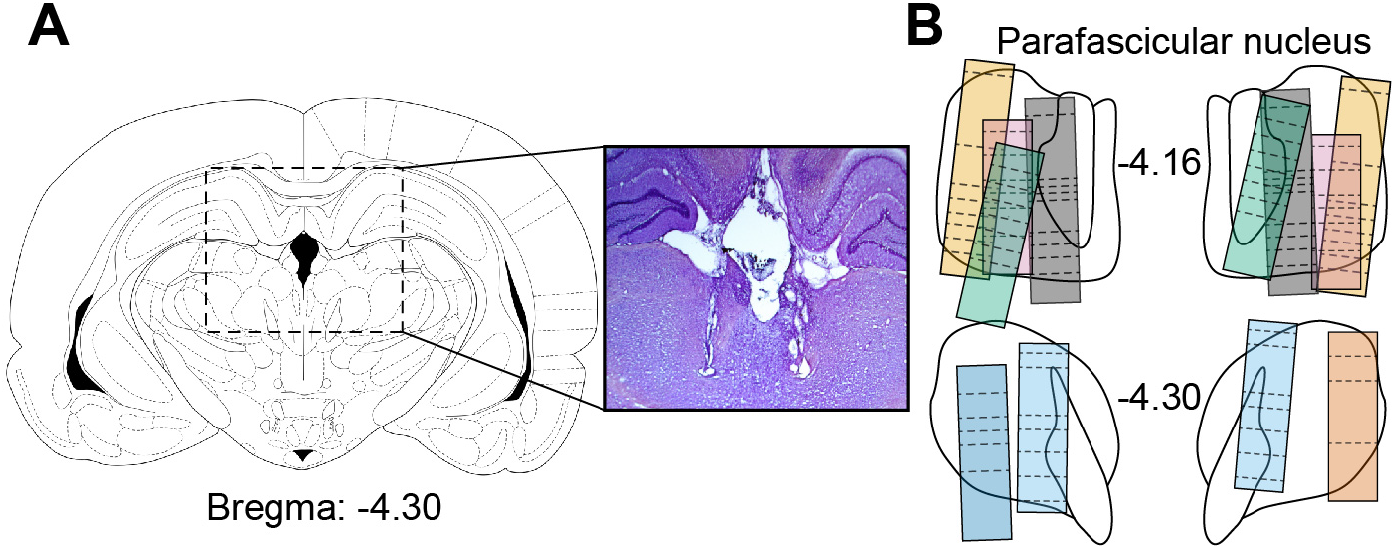
Histology and electrode path reconstruction. **A.** Schematic diagram of a rat brain coronal section and representative photomicrograph of two electrode bundle tracks located in the Pf. **B.** Histological reconstruction of the recording sites in the Pf. Boxes represent the approximate extent of the electrode bundles in the 7 animals (color coded). Dotted line corresponds to the depth recording sessions.

**Fig.3.**
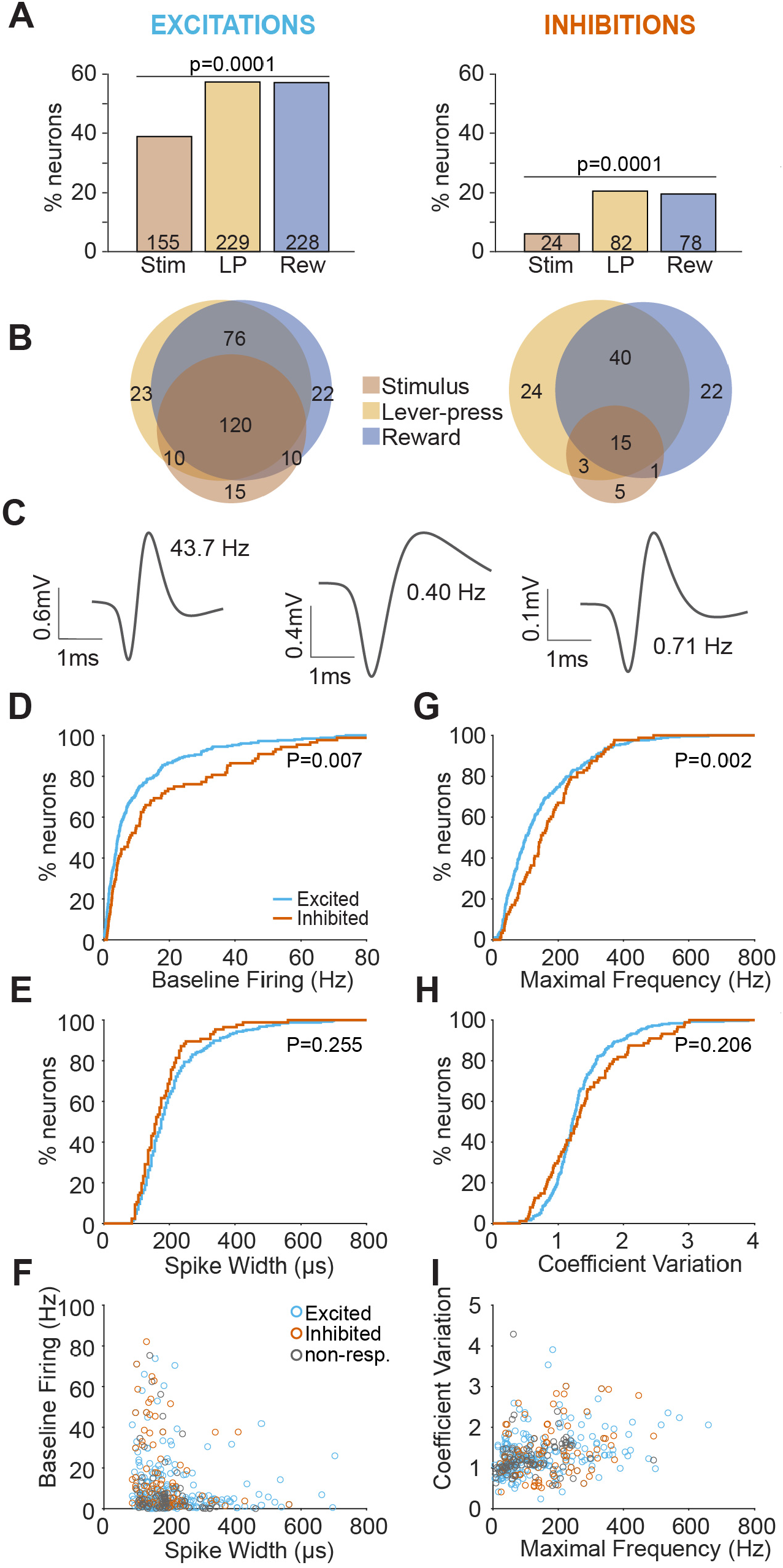
Electrophysiological characteristics of Pf neurons excited and inhibited by the different task events. **A.** Percentage of neurons excited (*left*) and inhibited (*right*) by the different events (stimulus, lever-press, reward). **B.** Venn diagrams showing the proportion of neurons excited (*left*) and inhibited (*right*) by task events. **C.** Example of three representative waveforms recorded in the Pf with their baseline firing rate. **D.** Cumulative percentage of baseline firing rate of task-excited (blue) and task-inhibited (red) neurons. **E.** Cumulative percentage of spike widths. **F.** Baseline firing rate plotted against spike width for individual neurons (gray dots indicate non-responsive neurons). **G.** Cumulative percentage of maximal frequencies **H.** Cumulative percentage of the coefficients of variation. **I.** Coefficient of variations plotted against the maximal frequencies for individual neurons.

We found that the majority of neurons were modulated by the different events with more excitations than inhibitions (254 and 88 neurons, respectively, χ^2^=141, P<0.0001). The excitations evoked by the stimulus were less frequent than those evoked by the lever-presses or the rewards (155, 229 and 228 neurons, respectively, χ^2^=27.9.0, P<0.0001, Fig. 3A) and the majority of neurons were excited to more than one event (Fig. 3B). Inhibitory responses evoked by stimuli were also less frequent than those evoked by lever-presses or rewards (24, 82 and 78 neurons, respectively, χ^2^=44.2, P<0.0001, Fig. 3A). The inhibitory responses to the different events showed less overlap than excitatory responses (χ^2^=37.2, P<0.0001, Fig. 3B).

Across the population, we observed neurons with different waveform profiles and firing properties during the baseline period (i.e. 10 s preceding stimulus onset, Fig. 3C). We analyzed separately the neurons excited or inhibited by task events. Event-excited neurons had an overall lower baseline firing rate (Kolmogorov-Smirnov test, KS=0.203, P=0.007, Fig. 3D) and a lower maximal frequency (KS=0.223, P=0.002, Fig. 3G) than event-inhibited neurons. The distributions of spike widths and coefficients of variations were similar between excited and inhibited neurons (KS=0.125, P=0.255; KS=0.130, P=0.206, respectively, Fig. 3E, H). When these variables were plotted against each other, event-excited and inhibited neurons strongly overlapped and it did not allow to isolate excited from inhibited neurons based on these different properties (Fig. 3F, I). We conducted similar analyses on excited and inhibited neurons separately for each event and obtained similar results (data not shown). Taken together, these results indicate that even if excited and inhibited neurons had different properties, it was insufficient to separate them.

### Pf neuronal responses to task events

We then analyzed the temporal dynamics and magnitudes of these different event-evoked responses that often lasted several seconds. But at the behavioral level, the stimulus, lever-presses and reward collections occur in close temporal proximity, often with smaller latencies than the duration of the neuronal responses. PSTHs constructed around each of these events can potentially be distorted by the presence of the other events that are correlated with each other in time. Thus, PSTHs do not allow one to accurately account for the neuronal responses to each individual event. To circumvent this issue, we used an iterative deconvolution method that takes advantage of the trial-to-trial variability in the temporal relationship between the different events to parse out the neuronal responses of each individual events from the PSTHs (Ghazizadeh et al., 2010). For each neuron, we deconvolved single-event responses by using the optimal number of iterations that had a cross-validation error lower than the PSTHs (5.2+/-0.2 iterations for the entire dataset, Fig. 4, 5).

**Fig.4.**
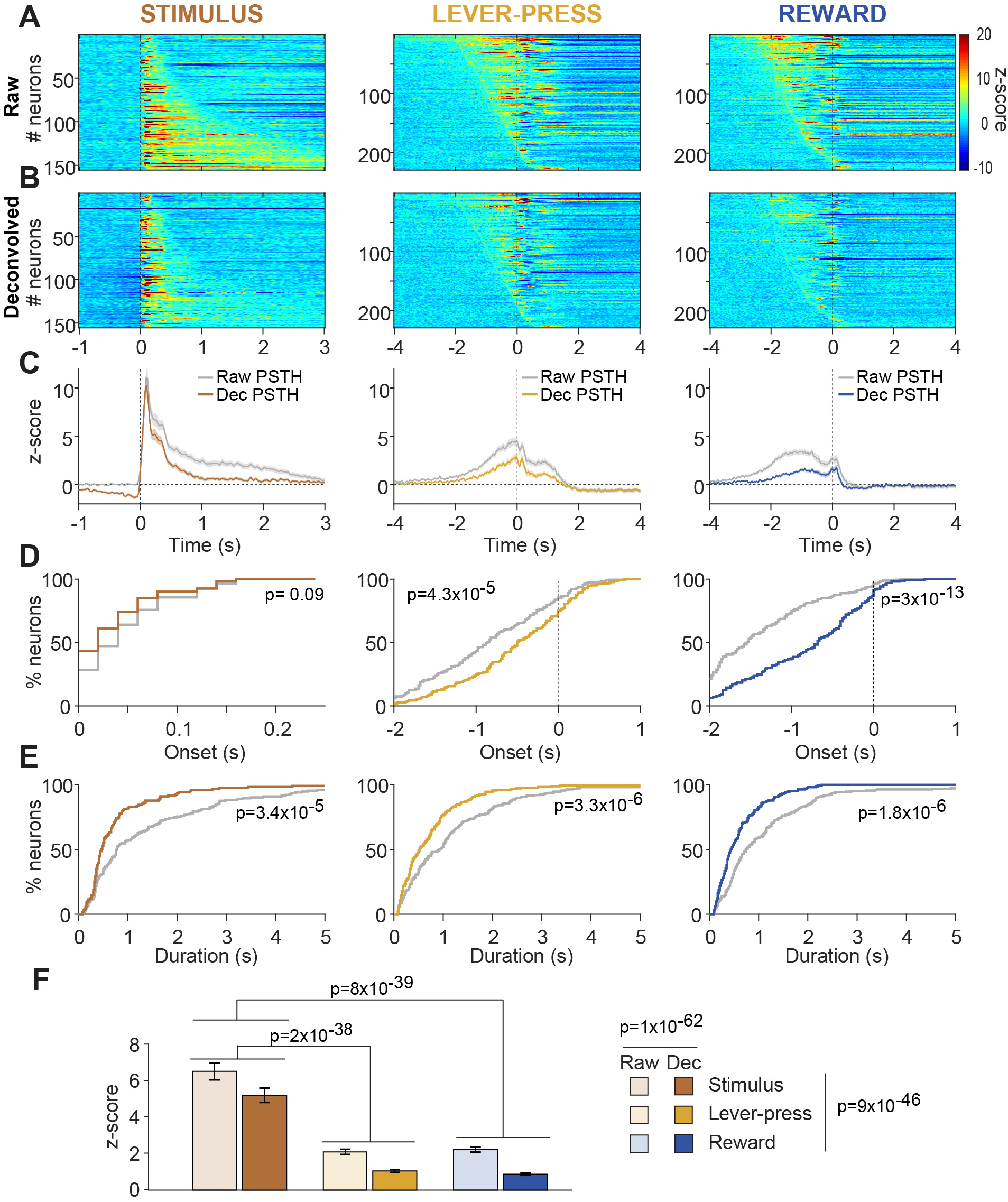
Pf neuronal excitations to task events. **A.** Heat-maps represent color-coded PSTHs showing neurons excited to the stimulus (*left*), lever-press (*middle*) and reward delivery (*right*). Each row represents the PSTH an individual neuron aligned to the event considered. Data are plotted with smoothed 20 ms-time-bins and neurons are sorted by excitation durations for stimuli responses and onset latencies for lever-press and reward responses. **B.** Heat-maps represent deconvolved PSTH of the same neurons represented in A. **C.** Average responses for the neurons shown in A and B. The gray traces correspond to raw PSTHs. Brown, orange and blue traces correspond to the deconvolved PSTHs to the stimulus, lever-press and reward delivery, respectively. **D.** Cumulative percentage of excitation onset latencies for raw and deconvolved responses. **E.** Cumulative percentage of excitations durations. **F.** Average z-scores of excitations.

**Fig.5.**
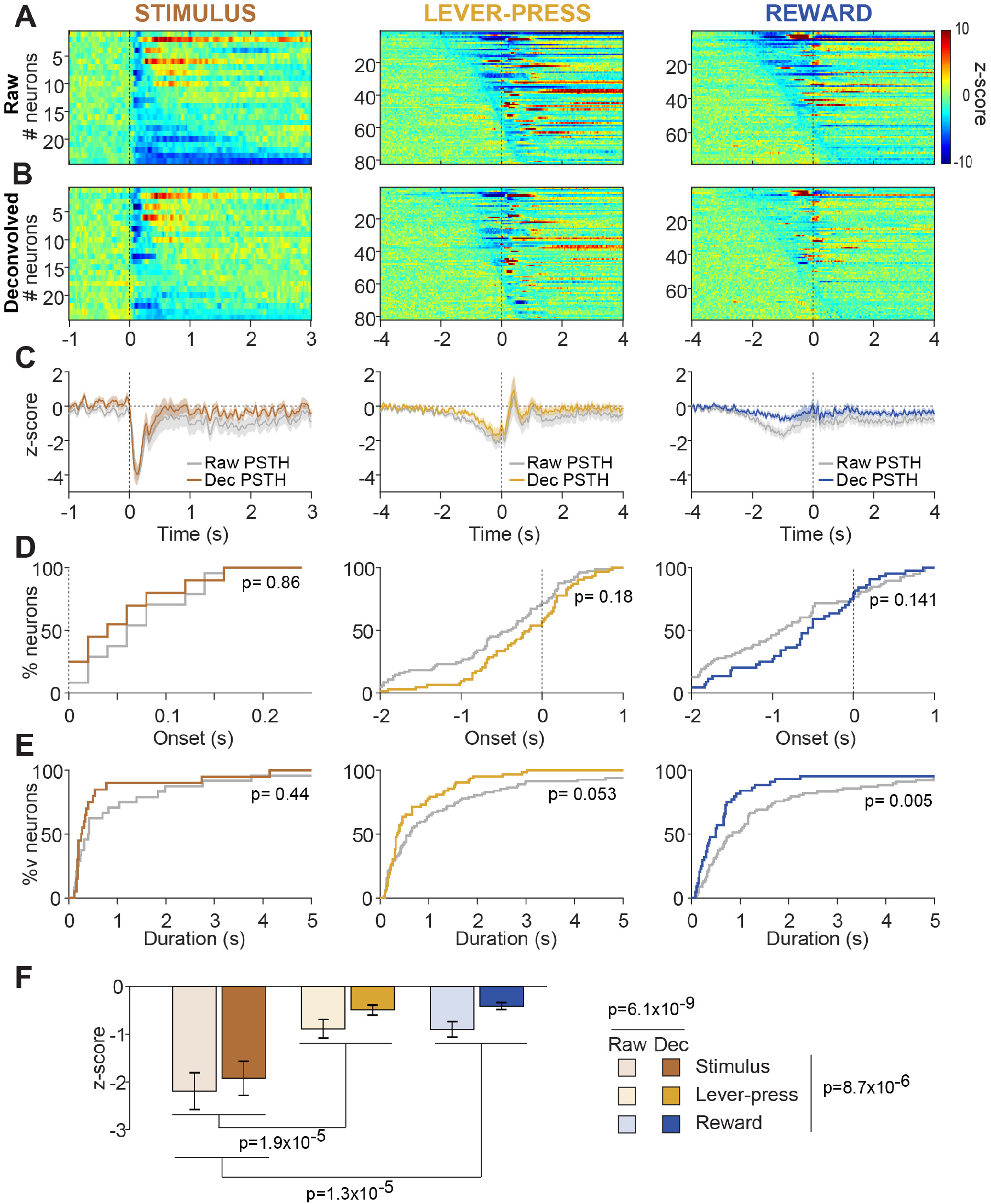
Pf neuronal inhibitions to task events. **A.** Heat-maps represent color-coded PSTHs showing neurons inhibited to the stimulus (*left*), lever-press (*middle*) and reward delivery (*right*). Each row represents the PSTH an individual neuron aligned to the event considered. Data are plotted with smoothed 20 ms-time-bins and neurons are sorted by inhibition durations for stimuli responses and onset latencies for lever-press and reward responses. **B.** Heat-maps represent deconvolved PSTH of the same neurons represented in A. **C.** Average responses for the neurons shown in A and B. The gray traces correspond to raw PSTHs. Brown, orange and blue traces correspond to the deconvolved PSTHs to the stimulus, lever-press and reward delivery, respectively. **D.** Cumulative percentage of inhibition onset latencies for raw and deconvolved responses. **E.** Cumulative percentage of inhibition durations. **F.** Average z-scores of inhibitions.

The analysis of raw PSTH aligned to the stimulus revealed that excitatory responses occurred at a stereotypical latency (58+/-4ms, Fig. 4A, C, D) with variable durations (1.57+/-0.17 s, Fig. 4A, C, E). The removal of the contribution of neighboring events by deconvolution had no effect on the onset latency distribution of stimulus-evoked excitations (KS=0.147, P=0.09, Fig. 4D) but strongly decreased their durations (KS=0.278, P=3.4×10^-5^, Fig. 4E).

With raw PSTH, most lever-press-evoked excitations emerged before the occurrence of the event (−0.830+/-0.05 s) and lasted up to 4 s (1.26+/-0.11 s). Reward-evoked excitations occurred even earlier (−1.36+/-0.04 s) and lasted in average 1.18+/-0.10s (Fig. 4A, C, D). Deconvolution significantly shifted lever-press- and reward-evoked excitations closer to the event (KS=0.226, P=4.3×10^-5^ and KS=0.382, P=3.0×10^-13^, respectively, Fig. 4D) and considerably shortened their durations (KS=0.25, P=3.3×10^-6^ and KS=0.26, P=1.8×10^-6^, respectively, Fig. 4E).

The analysis of the magnitude of evoked responses (Fig. 4F) revealed a strong influence of the event considered (2-way Repeated Measures ANOVA, Event effect, F_2,606_=123.6, P=9×10^-46^) and of the deconvolution (Deconvolution effect, F_1,606_= 355, P=1.1×10^-62^) but no significant interaction (Event x Deconvolution, F_2,602_=2.34, P=0.097). Stimuli-evoked excitations were ~3 times significantly larger than those evoked by lever-presses and rewards (Bonferroni test on Event effect, P=2×10^-38^ and P=8×10^-39^, respectively) but lever-press- and reward-evoked excitations did not differ from each other.

Inhibitions analyzed on raw PSTHs shared many similarities with excitations, but deconvolution had less effect, suggesting that these responses were more temporally associated with the behavioral events (Fig. 5). Stimulus-evoked inhibitory responses occurred after 85+/-9ms (non-significantly different from excitations, KS=0.264, P=0.09, Fig. 5A, B, D) and lasted 1.38+/-0.63 s (significantly shorter than excitations, KS=0.335, P=0.0014, Fig. 5E). Deconvolution had no effect on these measures (KS=0.175, P=0.85 and KS=0.25, P=0.44, respectively, Fig. 5B-E).

Inhibitions to lever-presses and rewards preceded the events for most neurons (−0.56+/-0.09 s and −0.82+/-0.10, respectively, Fig. 5A, C, D) and deconvolution had no effect (KS=0.18, P=0.18 and KS=0.21, P=0.14, respectively, Fig. 5B, C, D). We observed a strong trend toward a reduction of the duration of lever-press-evoked inhibitions by deconvolution (KS=0.22, P=0.053). The durations of reward-evoked inhibitions were reduced significantly (KS=0.312, P=0.006, Fig. 5E).

The magnitude of inhibitory responses (Fig. 5F) depended on the event considered (2-way Repeated Measures ANOVA, Event effect, F_2,181_=12.44, P=8.688×10^-6^) and the deconvolution (Deconvolution effect, F_1,181_=21.72, P=6.09×10^-6^) but we found no significant interaction (Event x Deconvolution, F_2,181_=0.53, P=0.585). Inhibitions to the stimulus were larger than those to the lever-presses and rewards (Bonferroni test on Event effect, P=1.9×10^-5^ and P=1.3×10^-5^, respectively) but lever-press- and reward-evoked excitations did not differ from each other.

Together, these results indicate that Pf neurons respond strongly to reward-predictive stimuli but that are also modulated in anticipations of the actions of lever-pressing and the collection of the rewards.

### Modulation of stimulus-evoked responses by the motivational state

Our data showed that Pf neurons are strongly activated by stimuli when the rats engage in reward-seeking. We then sought to decipher whether stimuli-evoked neuronal modulations depended on the motivational state. We took advantage of the fact that long sessions produce enough trials in which the rats engaged in reward-seeking in response to the stimulus (attended trials) and others in which they did not (unattended trials, Fig. 6). Visual inspection of the data revealed very brisk excitations leading us to use a higher time resolution (2 ms) to construct PSTHs and a shorter response duration requirement to detect excitations (4 ms). Because of the short latency of these responses that most certainly preceded locomotor onset to the stimulus (McGinty et al., 2013), we analyzed neuronal activity on raw and not deconvolved PSTHs. We observed a first population of 62 neurons (15.5%) with higher phasic activations in response to attended than unattended stimuli (paired t-test, t_61_=7.09, p=1.63×10^-9^, Fig. 6A, B). We named these neurons MOTIV + because their activation to the stimulus reflects the animal’s motivation to engage in the task. We also found a second population of 40 neurons (10%) that displayed an opposite pattern: they were more intensively excited to unattended than attended stimuli (MOTIV-neurons, paired t-test, t_39_=-4.89, P=1.77×10^-5^, Fig. 6A, B). The remaining neurons did not respond differently to the stimuli on attended and unattended trials.

**Fig.6.**
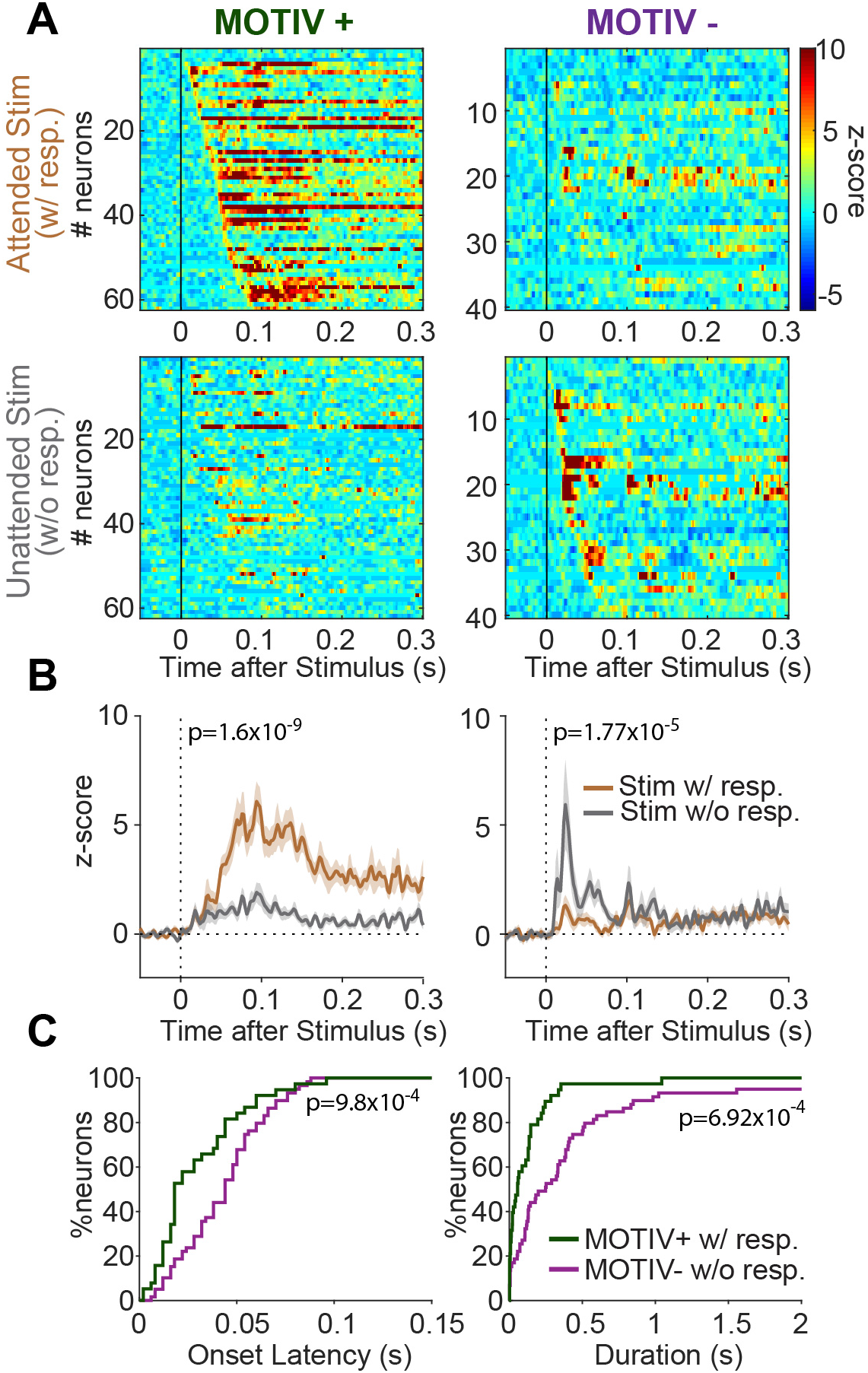
Pf neuronal excitations to the stimulus depend on whether the animal engages in rewardseeking. **A.** Heatmaps showing stimuli-evoked excitations of neurons on trials the animals engaged in reward-seeking (*top*) and those they did not (*bottom*) for MOTIV+ (*left*) *and MOTIV*-(*right*) neurons. Neurons are sorted by onset latencies. Raw PSTHs are plotted with 2 ms-time-bins. **B.** Average PSTHs for MOTIV + and MOTIV-neurons for attended (brown) and unattended (gray) stimuli. **C.** Cumulative percentage of excitation latencies (*left*) and durations (*right*) for MOTIV+ (green) and MOTIV-(purple) neurons.

The analysis of the response profile dynamics revealed that MOTIV-neurons were excited at a considerably shorter onset latency than MOTIV+ neurons (30.4+/-3.6 ms and 44.2+/-2.8 ms, respectively, KS=0.394, P=9.81×10^-4^, Fig. 6C). The excitations of MOTIV-neurons to unattended stimuli were also considerably shorter than those of MOTIV+ neurons to attended stimuli (117+/-29 ms and 436+/-87 ms, respectively, KS=0.403, P=6.92×10^-4^, Fig. 6C).

We conducted a similar analysis on stimuli-inhibited neurons on 20 ms-time-based PSTHs (Fig. 7) and found 21 MOTIV+ and 3 MOTIV-neurons. MOTIV+ neurons were more inhibited to attended than unattended stimuli (paired t-test, t_20_=-5.718, p=1.35×10^-5^). The onset latency to attended stimuli was 78+/-11 ms and lasted 535+/-132 ms. Given the low number of MOTIV-neurons, we did not conduct any comparison.

**Fig.7.**
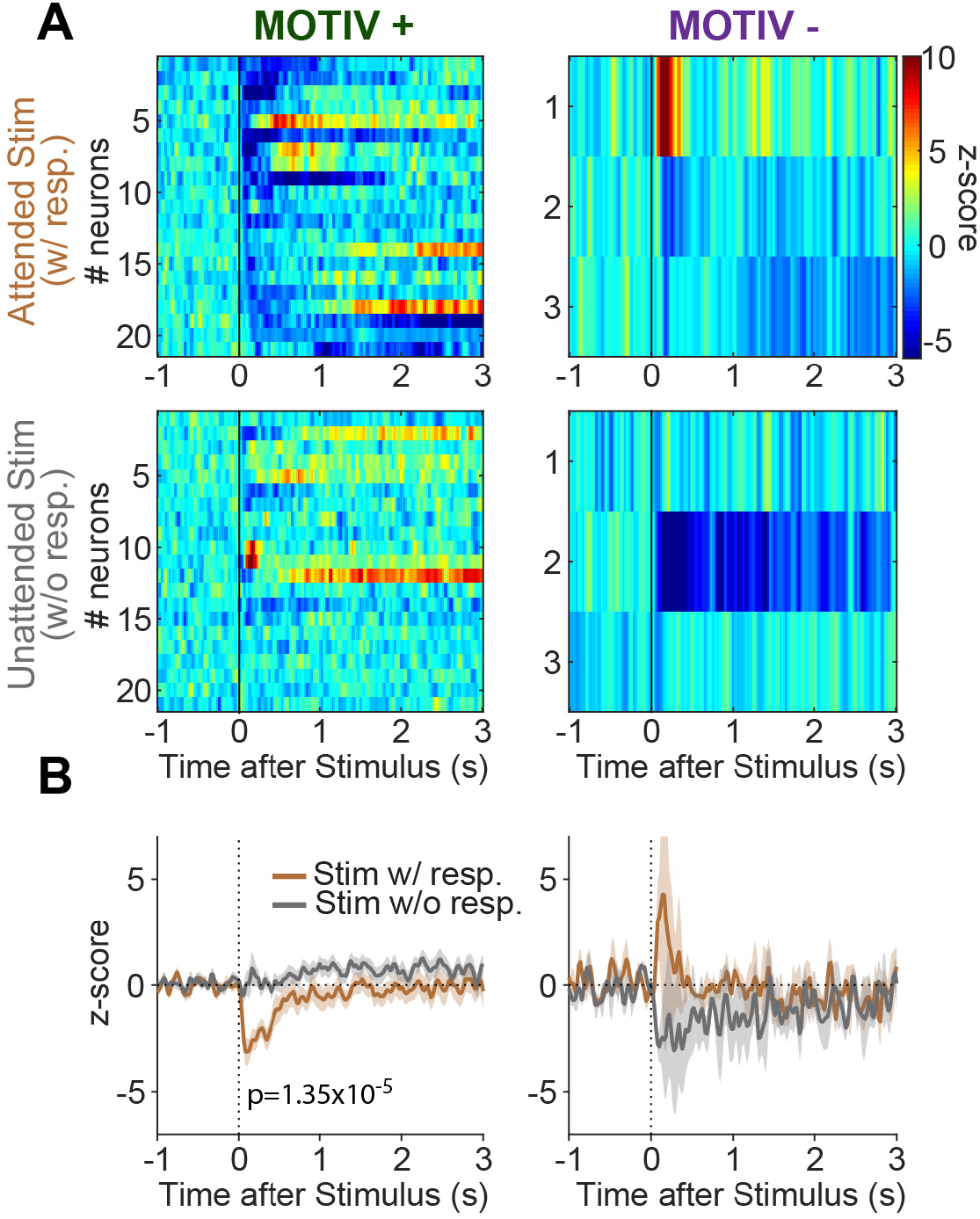
Pf neuronal inhibitions to the stimulus depends on whether the animal engages in reward-seeking. **A.** Heatmaps showing stimuli-evoked inhibitions of neurons on trials the animals engaged in reward-seeking (*top*) and those they did not (*bottom*) for MOTIV+ *(left) and MOTIV- (Iright)* neurons. Neurons are sorted by onset latencies. Raw PSTHs are plotted with 20 ms-time-bins. **B.** Average PSTHs for MOTIV + and MOTIV-neurons for attended (brown) and unattended (gray) stimuli.

## DISCUSSION

We sought to characterize the electrophysiological activity of Pf neurons in rats performing a stimulus-driven reward-seeking task. We show that most Pf neurons respond to the different task events with a higher proportion of excitations than inhibitions. The waveforms and discharge properties were not predictive of their excited or inhibited responses to task events. Stimuli evoked larger responses than the actions of lever-pressing and collecting the rewards. The most striking finding was that excitations to stimuli depended on whether the animal later engaged in reward-seeking.

### Basic electrophysiological properties of Pf neurons

Pf neurons have traditionally been identified as a homogeneous population of long-range projection glutamatergic neurons (Wilson et al., 1983; Mouroux and Féger, 1993; Smith et al., 2004). However, several studies have now revealed Pf subtypes with distinct morphological and electrophysiological signatures. Neurons with bushy dendritic trees or diffuse minimally branching dendrites show brief or long after-hyperpolarization, respectively leading to different abilities to sustain high frequency firing (Beatty et al., 2009; Mendez-Rodriguez et al., 2021). Furthermore, a recent study found that the mouse caudal Pf contains a small but significant proportions of GABA neurons intermingled with glutamate neurons (Chen et al., 2020). *In vitro*, GABA neurons displayed short action potential durations and higher discharge rates induced by current injections than glutamate neurons. To our knowledge, no studies have reported the presence of GABA neurons in rats. In light of these heterogeneous Pf neuron subtypes, we analyzed the basic electrophysiological properties of the neurons recorded extracellularly during the baseline period, when no task events were presented. The firing characteristics were assessed by measuring the basal firing rate and the maximal frequency reached as an index of their bursting abilities. Indeed, bushy neurons were reported to have higher maximal discharge rates and burstiness because of their shorter after-hyperpolarization (Beatty et al., 2009; Mendez-Rodriguez et al., 2021). We did find that all the parameters studied were dispersed but when plotted against each other, they strongly overlapped and we could not determine criteria to cluster different populations.

Because we found neurons excited and inhibited to different task events, we sought to determine whether these profiles mapped with their basic electrophysiological properties.

We did find that inhibited neurons had significantly higher basal firing rates and shorter spike widths. But here too, these differences were not sufficient to identify excited or inhibited neurons based on their intrinsic basal firing properties.

### Pf neurons are activated during the behavioral approach

We found a very high prevalence of neurons modulated by task events with a strong bias toward excitations. Stimuli clearly evoked the strongest responses, for both excited and inhibited neurons. This is certainly related to the temporal unpredictability of stimuli presentations and their association with rewards that have both been shown to potentiate the magnitude of excitations in the primate CM/Pf (Matsumoto et al., 2001; Minamimoto and Kimura, 2002). We found that stimuli-evoked responses occurred at a relatively constant latency with large variations in their duration. In many cases, we observed that excitations persisted until the rat lever-pressed. The long sessions introduced a large variability in the response latency within and between sessions that likely explained the variability in the durations of excitations. Deconvolution allows to isolate neuronal activity time-locked to the behavioral event considered by removing the contribution of neighboring events (Ghazizadeh et al., 2010; Ambroggi et al., 2011). The fact that deconvolution shortened stimulus- and action-evoked excitations suggests that the behavioral approach by itself activated Pf neurons. Interestingly, such pattern of anticipatory activity seems to be found in both primates and mice (Minamimoto et al., 2014; Díaz-Hernández et al., 2018). In a task where monkeys had to perform specific actions in response to instructive stimuli, the excitation of the classical long-latency facilitation (LLF) neurons recorded in CM appeared more strongly correlated with the timing of the required action than the instructive stimulus directing it (Minamimoto et al., 2014). Furthermore, a local inactivation of the CM with muscimol decreased the number of licks in a pavlovian task (Matsumoto et al., 2001), confirming the contribution of LLF activity on behavioral responding. In mice running a fixed-ratio 8 schedule task, where no explicit stimuli were presented, Pf neurons also displayed an excitation that preceded the first lever-press and that was shown to be necessary. Indeed, an optogenetic inhibition of striatum-projecting Pf neurons increased the latency to re-engage in a trial without affecting the repeated motor sequence of lever-pressing (Díaz-Hernández et al., 2018). Altogether, these results indicate that the excitation of Pf neurons during the behavioral approach is causal.

### Influence of the motivational state on Pf responses to incentive stimuli

The use of long sessions allowed to analyze Pf neuronal activity in response to stimuli predicting the same rewards while rats were in different motivational states. We compared the trials to which the animal attended to the stimulus by engaging in the behavioral response (and thus obtaining the reward) and those they did not. The absence of engagement on unattended trials was unlikely due to a failure in perceiving the stimulus that was highly salient (85 dB white noise coupled to the visual extension of the lever) and long lasting (up to 10 s). The strongest evidence against this hypothesis is the fact that we identified a population of neurons that was more excited on unattended trials, indicative of an active process taking place in these situations. Most likely, the absence of responding was caused by the fact that at certain times, animals in these long sessions valued reward-seeking less than other activities (e.g. grooming, exploring, resting).

MOTIV+ neurons exhibited either excitations or inhibitions to attended stimuli in the first 100ms of their presentations. In a similar task, locomotion onset has been reported to start ~250ms after a reward-predictive stimulus, indicating that the initial component of the responses of MOTIV+ neurons was not driven by movements themselves (McGinty et al., 2013) but could participate in the initiation process. It seems unlikely that MOTIV+ neurons are homolog to primate LLF neurons for two reasons. First, LLF neurons usually have a biphasic response with an inhibition preceding the excitation (Matsumoto et al., 2001) which we did not observe in MOTIV+ neurons. Second, the amplitude of the excitatory component of LLF neurons is inversely correlated with the reaction time on a trial-to-trial basis (Minamimoto et al., 2014). The stronger response to attended stimuli compared to unattended stimuli is diametrically opposed to this observation. Thus, the MOTIV+ profile reported in this study does not seem to match LLF activity or any other reported in primates that we are aware of.

MOTIV-neurons had a stereotyped response with excitations starting as early as 6 ms after stimulus onset and lasting less than 100ms. Yet, these brisk responses were associated with the absence of behavioral responding in the next 10 s. These data provide further evidence that the Pf carries an important attentional function (Minamimoto and Kimura, 2002; Minamimoto et al., 2009; Redgrave et al., 2011; Smith et al., 2011) by gathering low level sensory information that may arise from the deep layers of the superior colliculi and/or the pedunculopontine nucleus (Paré et al., 1988; Krauthamer et al., 1992; Krout et al., 2001).

MOTIV-neurons shared properties with short-latency facilitation neurons (SLF) recorded in the primate Pf (Matsumoto et al., 2001). But importantly, the early and transient firing of MOTIV-neurons manifested when the rat did not attend to stimuli indicates that their attention was not directed toward them. To the contrary, Pf recordings and inactivations in monkeys during a countermanding task provided evidence that SLF neurons participate in the direction of attention toward stimuli (Minamimoto and Kimura, 2002). This apparent discrepancy could relate to the modality used in these studies (auditory and visual in our study and purely visual in the primate study) or even different functions carried by Pf neurons in different species. Another intriguing possibility lies in the characteristics of the stimuli used. Minamimoto and Kimura (2002) presented temporally predictable stimuli that provided instructions to the monkeys about the direction of the saccade to be rewarded. In our study, we used stimuli that were presented unexpectedly and incentivized the rats to switch from their current activity to reward-seeking by engaging in actions during a 10 s-time window. These actions strongly differed depending on the location of the rats at the time of occurrence of these stimuli. Thus, the Pf could subserve different roles for instructive and incentive stimuli as we previously showed for NAc neurons (Sicre et al., 2020).

The excitations of MOTIV-neurons could participate in the process of not engaging in actions in response to the incentive stimulus through the dense Pf projection to the striatal complex (Van der Werf et al., 2002). As opposed to other thalamic nuclei, the Pf preferentially synapses on cholinergic interneurons (CINs) (Lapper and Bolam, 1992; Sidibé and Smith, 1999; Raju et al., 2006; Doig et al., 2014) that exert a strong inhibitory control on the activity of medium spiny projection neurons (MSNs) *in vivo* (Witten et al., 2010) through different GABA interneuron subtypes (English et al., 2012; Assous et al., 2017; Assous and Tepper, 2018). We recently reported that NAc Core CINs were also more active in response to unattended than attended incentive stimuli (Sicre et al., 2020). The short latencies observed in CINs suggest that they could be driven by MOTIV-Pf neurons.

Excitations of NAc Core MSNs evoked by incentive stimuli are necessary for rats to engage in action (Ambroggi et al., 2011) and depend on ventral tegmental area (Yun et al., 2004), basolateral amygdala (Ambroggi et al., 2008) and paraventricular (Meffre et al., 2019) inputs. Together, our recent work suggests a circuit able to repress the activation of NAc Core MSNs to incentive stimuli by these inputs when the animal is not willing to engage in action through the activation of CINs driven by Pf MOTIV-neurons. This circuit may allow to dynamically filter which predictive information is able to control NAc Core MSNs and thus suggest an attentional process directed toward the motivational system.

## ABBREVIATIONS

CIN: Cholinergic interneurons
CM: Center median nucleus of the thalamus
FR: Fixed-ratio
LLF: long-latency facilitation
MSN: Medium spiny neurons
NAc: Nucleus accumbens
Pf: Parafascicular nucleus of the thalamus
PSTH: Peri-stimulus time histograms
SLF: short-latency facilitation

## CONTRIBUTIONS

MS conducted experiments, analyzed data, prepared figures and wrote the initial draft of the manuscript. JM supervised MS, participated to experiments and provided critical comments to the manuscript. FA designed research, analyzed data, prepared figures and edited the manuscript.

## ACKNOWLEDGEMENTS

The authors would like to thank Bruno Poucet for his support, Didier Louber and Dany Paleressompoulle for technical assistance and Simon Moré for information technology assistance.

